# Mechanism of Antiviral Activity of Nitazoxanide against the Influenza Virus: Effect of Tizoxanide on AdenosineTriphosphate in Influenza-virus Infected Madin Darby Canine Kidney Cells

**DOI:** 10.1101/2021.07.30.454324

**Authors:** J.F. Rossignol, A.S.L. Tijsma, C.A. van Baalen

## Abstract

**Background:** Nitazoxanide (NTZ) is a broad-spectrum antiviral undergoing clinical development for treating influenza and other viral respiratory infections such as those caused by rhinovirus/enterovirus and coronavirus including the emerging SARS-CoV-2.

**Methods:** Nitazoxanide is a mild uncoupler of oxidative phosphorylation, which is modulating the ATP production in cells. ATP is an essential component of viral replication, and we have evaluated the effect of tizoxanide (TIZ), the active circulating metabolite of NTZ, on ATP in Madin-Darby canine kidney (MDCK) cells and in MDCK cells infected with influenza A and B viruses.

**Results:** TIZ decreased cellular ATP in a dose-dependent manner in MDCK cells and in MDCK cells infected with influenza A and B viruses. Maximum inhibition of ATP in influenza infected or uninfected MDCK cells reached up to 45% after 6 and 24 hours of exposure to 100µM TIZ. The decrease in cellular ATP did not affect cell viability and was reversible after eliminating TIZ from the culture.

**Conclusion:** The concentrations of TIZ required to decrease cellular ATP levels were similar to those reported to inhibit replication of influenza A and B viruses in our laboratory. A decrease in ATP triggers activation of AMP-activated protein kinase, which is known to suppress the secretion of pro-inflammatory cytokines. Additional studies are warranted to evaluate the effect of TIZ on mitochondrial function.

## 1. Introduction

Nitazoxanide (NTZ) is a broad-spectrum antiviral agent undergoing clinical development for treating influenza and other viral respiratory diseases including COVID-19^1-3^. It is a mild uncoupler of oxidative phosphorylation.^4^ Mechanistic studies have shown that NTZ and its active circulating metabolite, tizoxanide (TIZ), block maturation of the influenza hemagglutinin glycoprotein at the post-translational stage.^5^ The broad-spectrum antiviral activity of NTZ and the inability to select for resistance suggest a host target, but a specific target has not yet been clearly identified.^6^ In an effort to understand the mechanism of antiviral activity of TIZ better, we evaluated the effect of TIZ on ATP in MDCK cells and in MDCK cells infected with influenza A and B viruses.

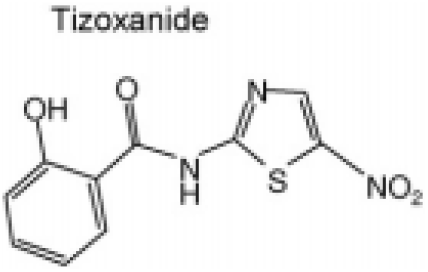

## 2. Material and Methods

### 2.1 Material Compounds

TIZ was provided by Romark (Tampa, Florida USA). Working stocks of TIZ dissolved in DMSO were prepared and stored at -80°C until used.

#### Cell cultures

Madin-Darby canine kidney (MDCK) cells were passaged in flasks and seeded in 96-well plates. Before inoculation, cells were cultured in MDCK medium and after inoculation in infection medium containing N-tosyl-L-phenylalanyl chloromethyl ketone (TPCK)-treated trypsin (3µg/mL; Sigma, T1426).

### 2.2 Methods

#### Measurement of ATP

ATP measurements were performed using the CellTiter-Glo Luminescent Cell Viability Assay (Promega, Madison, Wisconsin USA) according to the manufacturer’s instructions, with the exception that cell culture medium was aspirated prior to adding 100 µL of ATP reagent per well. Opaque walled 96-well plates were used.

#### Measurement of dehydrogenase activity

Dehydrogenase activity was measured using the Cell Counting Kit-8 (Sigma-Aldrich). Twenty-µL WST-8 reagent was added to each well of 96-well plates already containing 200 µL cell culture medium per well and incubated for 1-2 hours at 37 °C. After incubation, absorbance was measured at 450 nm.

#### Experiments in uninfected MDCK cells

Multiple identical plates with MDCK cells were prepared. Cellular ATP levels and dehydrogenase activity were measured after 6 and 24 hours of exposure to various TIZ concentrations (0 to 100µM). Nuclei of MDCK cells were stained with DAPI (Life technologies) after treatment with TIZ for 6 or 24 hours. Images were taken using a CTL ImmunoSpot S6 Analyzer and in a representative area of the well, the cell nuclei were quantified using Immunospot software. Using identical replicate plates, the compound was removed; cells were incubated again in fresh cell culture medium (without TIZ) for 24 hours after which cell nuclei were quantified again.

#### Experiments in influenza virus-infected MDCK cells

Multiple identical 96-well plates with MDCK cells were (mock) infected with two influenza A and two influenza B strains for one hour at low and high MOI. After this period, the inoculum was aspirated and replaced by medium containing various TIZ concentrations (0, 10, 30, 100µM). The cell cultures were incubated for an additional five hours (single cycle infection, high MOI) or 23 hours (multi-cycle infection, low MOI) after which cellular ATP levels were determined. In addition, cells were immunostained for influenza (for influenza A: HB65, EVL; for influenza B: MAB8661, Millipore). Images were taken using a CTL ImmunoSpot S6 Analyzer and the percentage well area covered (WAC) by influenza-positive cells was quantified using Immunospot software. The virus input was checked by back titration.

## 3. Results

### Uninfected MDCK cells

At six hours post-TIZ treatment, a concentration-dependent decrease of cellular ATP levels was observed, starting at 12.5 µM TIZ and higher with a maximum ATP decrease of 45% at 100µM TIZ. After 24 hours of treatment with TIZ, the maximum reduction was similar (44% at 100µM); however, reduction of ATP was observed at lower concentrations of TIZ (6.25µM) as compared to 6 hours.

To assess if TIZ affected cell viability, we also measured dehydrogenase activity because it is often used as an ATP-independent measure for cell viability. After 6 hours of exposure to TIZ, dehydrogenase activity was at the level of untreated control cells, although at high concentrations of TIZ (≥50µM), the activity increased somewhat (Fig. 1A). After 24 hours of TIZ treatment, the dehydrogenase activity decreased at higher (≥50µM) TIZ concentrations (Fig. 1B).

**Fig. 1.**
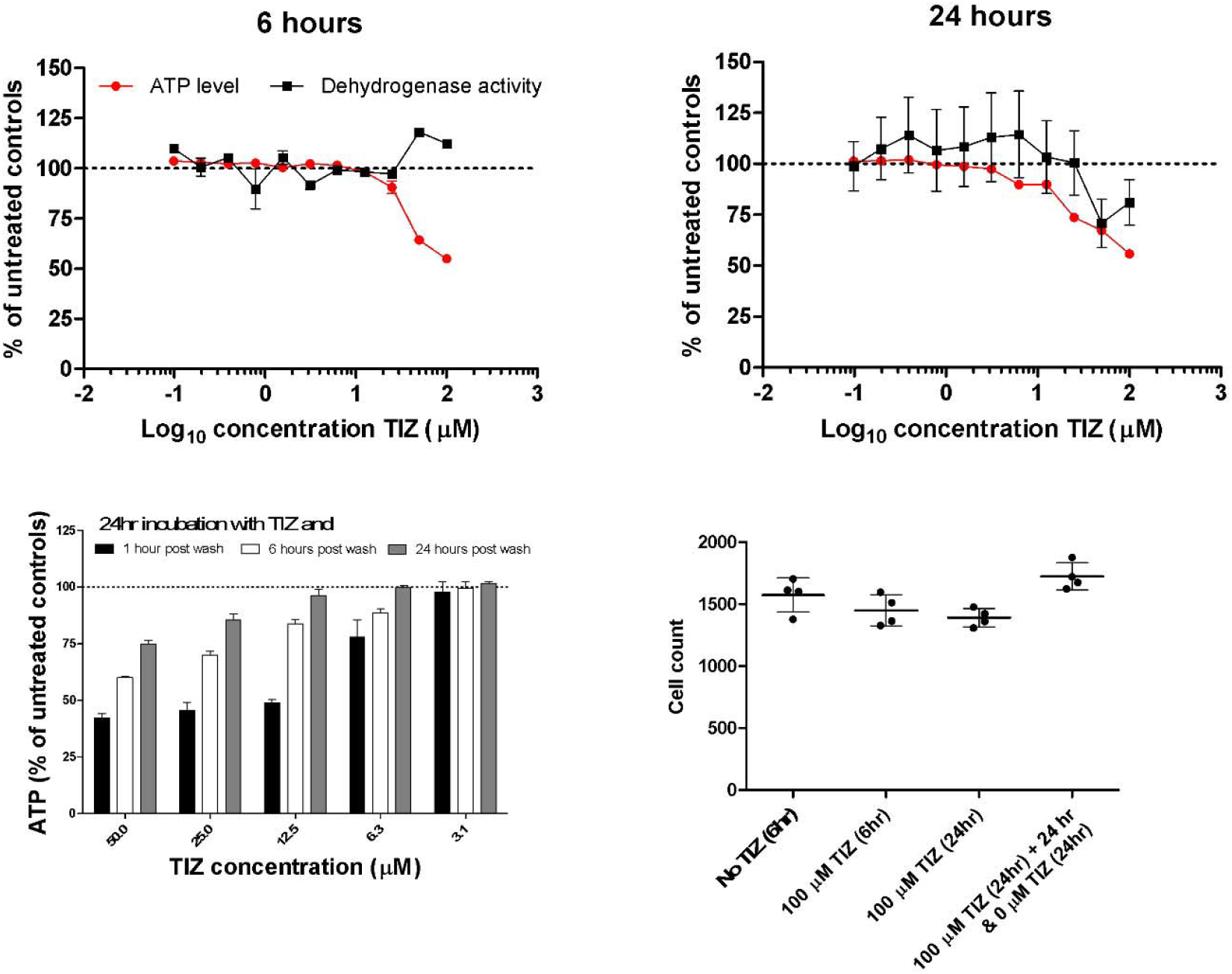
Effect of TIZ on ATP levels, dehydrogenase activity and cell count in uninfected MDCK cells. Effect of TIZ on ATP levels and dehydrogenase activity after 6 (A) and 24 (B) hours of incubation with the compound. Effect of TIZ on ATP levels were also assessed after 24 hours of incubation with TIZ after which the medium was refreshed (not containing TIZ) and subsequent incubation for 1, 6 and 24 hours (C). The effect of TIZ on the number of attached cells was determined by counting DAPI stained cell nuclei after various incubation times with 100 µM of TIZ (D). Each datapoint is the mean ± SD of 4 replicates.

To evaluate whether the effect of TIZ on ATP is transient or enduring, MDCK cells were treated with various concentrations of TIZ for 24 hours. After this period, the cells were washed once with PBS to remove TIZ, and fresh cell culture medium (not containing TIZ) was added to the cells. After a subsequent incubation period of 1, 6, and 24 hours, the cellular ATP levels were determined (Fig. 1C). In this experiment, cellular ATP levels recovered with increasing incubation time. At concentrations of 12.5µM and lower, ATP levels recovered to the level of untreated control cells. However, at higher concentrations (≥25µM), full recovery did not occur within the timeframe of 24 hours.

To exclude the possibility that reduced ATP levels and dehydrogenase activity are due to a reduced number of cells in the wells (due to detachment, apoptosis and necrosis for instance), MDCK cells nuclei were stained with DAPI and quantified once TIZ treatment was completed (Fig. 1D). The number of cells did not significantly decrease after treatment with 100µM TIZ for 6 or 24 hours. Moreover, when cells were incubated with 100µM TIZ, after which the compound was removed and incubated again in fresh cell culture medium for 24 hours, the number of nuclei increased, most likely due to cell proliferation.

### Influenza virus-infected MDCK cells

We then measured ATP levels in MDCK cells infected with two influenza A virus strains and two influenza B virus strains using both single cycle (6 hours, multiplicity of infection (MOI) > 1) and multicycle (24 hours, MOI ≤ 0.01) infection. For both single- and multicycle infection (Fig. 2A and 2B), TIZ reduced ATP in influenza-infected cells in a concentration-dependent manner, similarly to its effect in uninfected cells. For the single cycle infection (Fig 2C), back titration showed that the MOI used in the assay varied between 1.2 and 20.4. Virus replication was assessed by influenza-specific immunostaining of the cell monolayer. The well area covered by the immunostaining was calculated and is a measure of virus replication. The maximum reduction of well area covered by influenza virus spots was 3.7-fold when cell cultures were treated with 100µM TIZ. At 30 and 10 µM, the reduction was at most 1.5-fold.

**Fig. 2.**
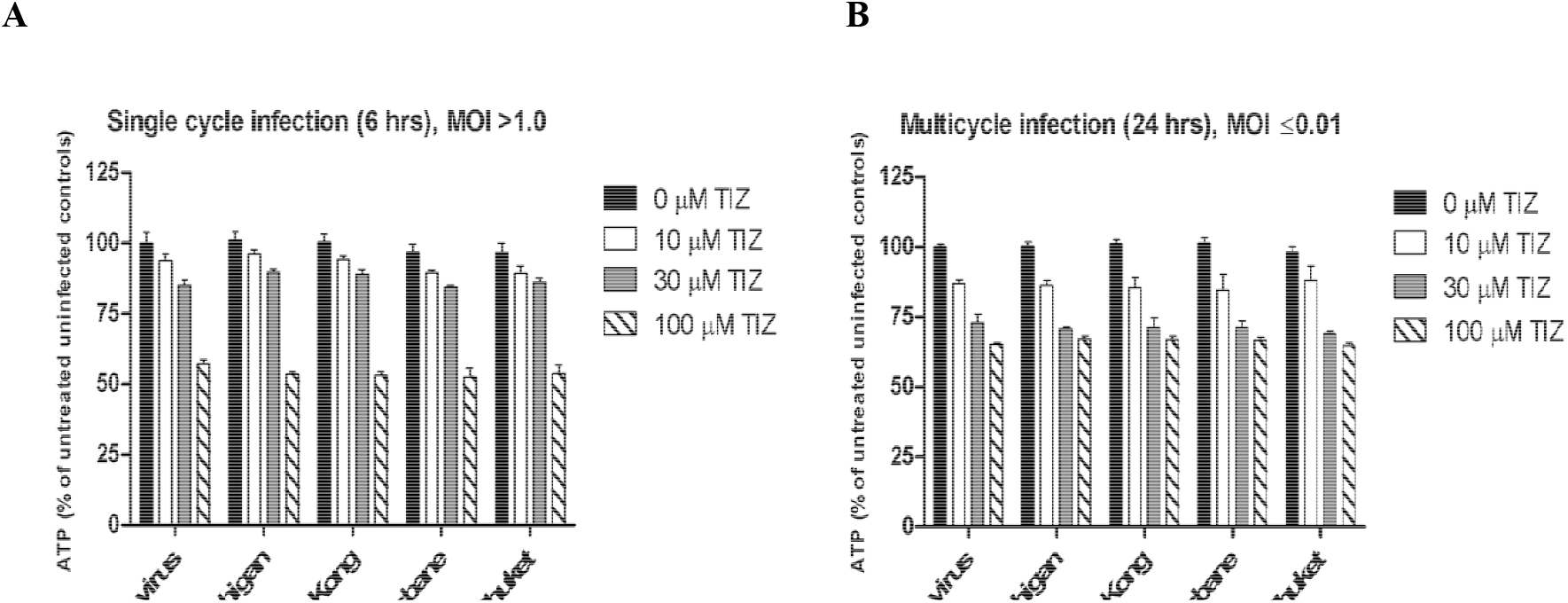
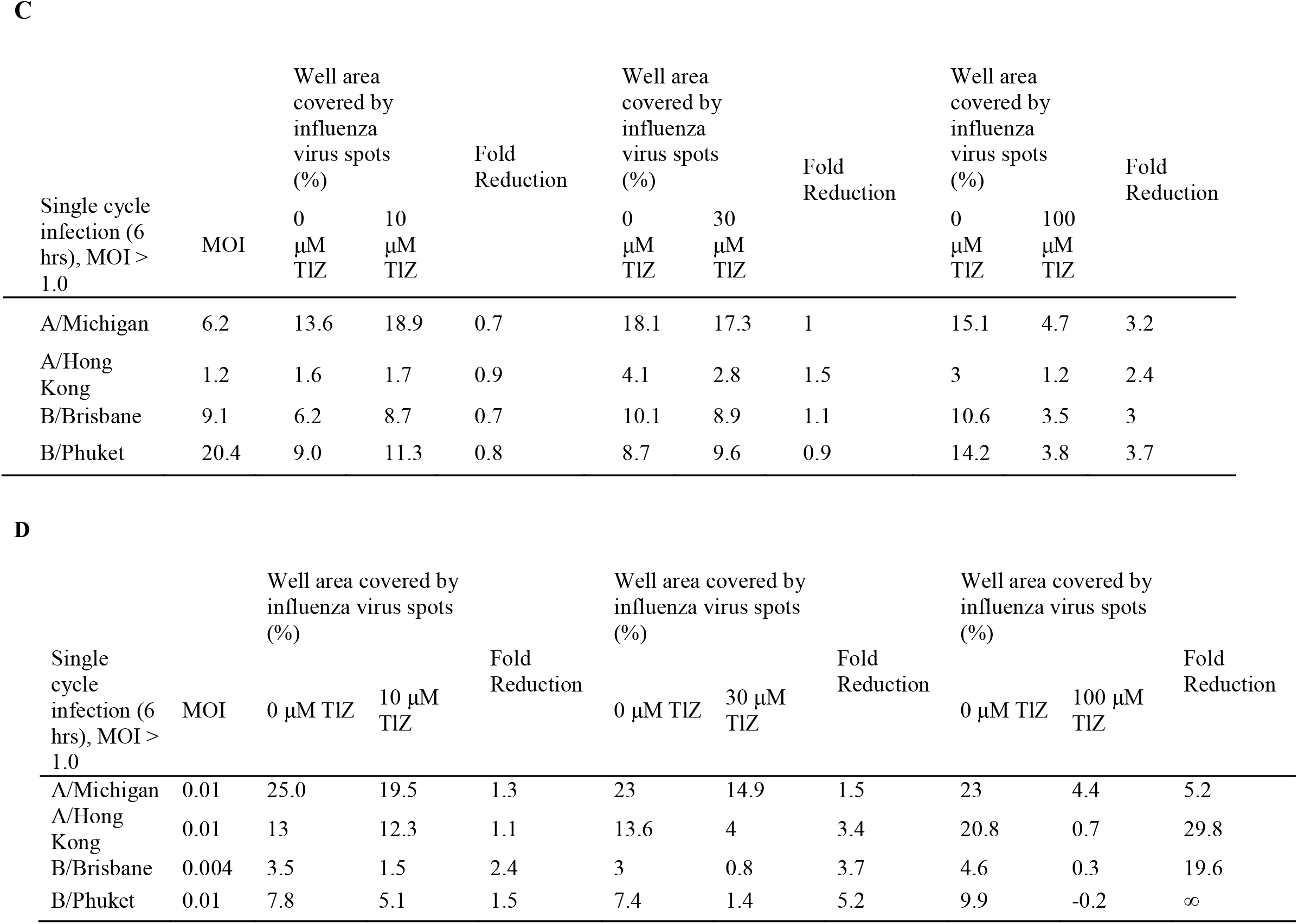
TIZ decreases ATP in influenza virus-infected MDCK cells during single cycle infection (A) and multi-cycle infection (B). Data depicted in the graph are the means of at least 4 determinations, error bars represent the SD. Fold reduction in well area covered (WAC) by influenza virus spots (%) in TIZ treated cultures compared t untreated cultures in single cycle (C) and multicycle infection (D). Percentages shown are the means of at least 4 replicates. Background %WAC (uninfected cell controls) is subtracted to calculate the specific %WAC of eac culture condition, shown here.

For the multicycle infection (Fig 2D), back titration results showed that the MOI used in the assay varied between 0.004 and 0.01. Full inhibition of virus replication was observed for B/Phuket when incubated with 100 µM TIZ. The negative well area covered (WAC) for this culture condition (−0.2) i because the background WAC was slightly higher than the specific WAC of this culture condition; hence, no fold reduction could be calculated, which was denoted with “∞”. For the other viruses, treated with 100 µM TIZ, the fold reduction varied between 5.2 and 29.8-fold. At 30 and 10 µM TIZ, the reduction was at most 5.2-fold. It has to be noted that the fold reduction for A/Michigan at 100 and 30 µM TIZ is likely to be larger than could be calculated since the wells not treated with TIZ were completely positive for influenza NP, meaning that uninhibited virus propagation was capped by the well area.

## 4. Discussion

These data show that TIZ decreases cellular ATP in a dose-dependent manner in MDCK cells and in MDCK cells infected with influenza A and B viruses. Maximum inhibition of ATP in influenza infected or uninfected MDCK cells reached up to 45% after 6 and 24 hours of exposure to 100µM TIZ. The decrease in cellular ATP did not affect cell viability and was reversible after removing TIZ from the culture. The concentrations of TIZ required to decrease cellular ATP levels were similar to those reported to inhibit replication of influenza A and B viruses in our laboratory.

Studies of a number of different viruses have shown that viral replication is ATP-dependent. Braakman et al. first described the role of ATP in the formation of disulphide bonds during the maturation and the protein folding of the influenza hemagglutinin in the endoplasmic reticulum in 1991.^7^ The following year, 1992, Braakman et al further studied the effects of ATP on viral replications.^8^ Their results were consistent with our results on the effect of NTZ on the maturation of influenza glycoprotein in the endoplasmic reticulum.^5^ It was later shown that ATP is regulating the assembly and the transport of vesicular stomatitis virus G protein trimers^9^ and that an increased ATP generation in the host cells was required for an efficient vaccinia virus production.^10^ Finally, Mirazimi & Svensson showed that ATP was an essential component for correct folding and disulphide bond formation in rotavirus.^11^

NTZ and TIZ are modulators of mitochondrial activity by uncoupling oxidative phosphorylation.^4^ NTZ targets the maturation of key viral proteins in the endoplasmic reticulum, hemagglutinin (influenza), F-protein, (RSV & parainfluenza) and N-protein (coronavirus).^5,12-13^ Effectively, we have been able to show that NTZ blocks the proper folding of proteins of two paramyxoviridae, the respiratory syncytial virus (RSV) or the Sendai virus (SeV) in blocking the effect of disulfide isomerase ERp57.^12^

An indirect effect of ATP inhibition by targeting mitochondrial activity is triggering the phosphorylation of the AMP-activated protein kinase (AMPK) suppressing the secretion of pro-inflammatory cytokines^14-16^ while in a few cases such as metformin the activation of AMPK is not affected by the ATP level in the cells.^17^

Anti-cytokines activity of TIZ was demonstrated in cell bioassays against seven pro-inflammatory cytokines, TNF-alpha, Il-2, IL-4, IL-5, IL-6, IL-8 and IL-10 with IC_50_s ranging from 0.67 to 2.65 µg/mL. Hong et all have also investigated the effect of NTZ on pro-inflammatory cytokines.^18^ The drug suppresses lipopolysaccharide (LPS)-induced production of IL-6 from RAW 264.7 cells and mouse peritoneal macrophages with 50% inhibitory concentrations (IC_50_s) of 1.54 µM and 0.17 µM, respectively. NTZ also inhibited the LPS-induced expression of IL-6 mRNA in RAW 264.7 cells. *In vivo* the drug was orally administered at a dose of 100 mg/kg to mice 2 hours before a 1 mL intraperitoneal injection of 4% thioglycollate (TG). Six hours after TG injection, plasma IL-6 levels were markedly lower (by 90%) than the levels in vehicle-treated mice. These data suggest that nitazoxanide could be a promising drug against various diseases associated with overproduction of IL-6.^18^

## 5. Conclusion

The antiviral mechanism of action of NTZ/TIZ is related to the energy metabolism process in cells. The modulating effect of the drug on mitochondria results in a modest decrease of the ATP concentration in influenza infected and in non-infected cells. This reduction does not affect the viability of the cells but prevents the maturation of viral proteins in the endoplasmic reticulum. In addition, a decrease of ATP is activating the AMP-activated protein kinase (AMPK) responsible for blocking the secretion of several pro-inflammatory cytokines. In effect, cell-based assays showed that concentrations of tizoxanide prevent the secretion of TNF-alpha, Il-1, IL-2, IL-3, IL-5, IL-8, IL-10 with IC_50_ from 0.67 to 2.65 µg/mL. Effects against IL-6 have also been reported in cell assays and *in vivo*. However, this anti-cytokine activity may not be clinically significant and needs to be proven in patients infected with a viral illness.

## Acknowledgment

Jean-François Rossignol is the inventor and the developer of Nitazoxanide. He holds several patents on the drug and its applications. He also is the co-founder of Romark Laboratories, L.C. a stockholder and an employee of the company. Carel van Baalen and Aloys Tijsma are employees of Viroclinics Biosciences B.V. in Rotterdam, The Netherlands.

## Authors Contribution

Jean-François Rossignol designed the aforementioned experiment, and he is the primary author of this publication. Carl van Baalen and Aloys Tijsma performed the experimental research under a contract with Romark Laboratories L.C.

